# Adolescent predatory stress increases peripheral TNFα response to LPS regardless of sex

**DOI:** 10.1101/2021.01.25.427666

**Authors:** Samya K. Dyer, Kimaya R. Council, Susie Turkson, Gladys A. Shaw, Gretchen N. Neigh

## Abstract

Chronic adolescent stress has been shown to cause increased adverse effects in response to an immune stressor later in life including the development of anxiety and PTSD. Following stress, the production of inflammatory cytokines including IL-1β and TNFα is expected to increase in response to an immune challenge such as LPS. Male and female mice underwent fifteen consecutive days of predatory stress. Chronic adolescent stress was shown to increase both anxiety-like behavior (F_(1, 28)_ = 19.82, p = 0.0001) and TNFα levels within the periphery (F_(1, 19)_ = 4.748, p = 0.0421).

## Introduction

Chronic stress is a major risk factor in the development of anxiety disorders, including post-traumatic stress disorder (PTSD) (Brown et al., 2014). Typically, our bodies quickly recover from bouts of acute stress, both positive and negative. However, chronic stress can cause irreparable damage within the brain’s neural circuitry. The likelihood of anxiety disorder development increases concurrently with trauma load across one’s lifespan (McEwen, 2007). In addition, the effects of chronic stress become even more detrimental when the stress occurs during adolescence (Blakemore et al., 2010). As a time of intense neuronal and endocrine development, adolescence is considered an especially vulnerable time period (Spear, 2000), with repeated stress inducing increases in mental health disorders in humans (Andersen & Teicher, 2008) and anxiety-like behavior in rodents (Bourke and Neigh, 2011) that persists well into adulthood (Burgado et al., 2014).

Chronic stress not only negatively impacts neural development caused by dysregulation within the Hypothalamic-Pituitary-Adrenal (HPA)-axis leading to alterations in the neurobiological substrates related to anxiety disorders (Neigh et al., 2013), chronic adolescent stress also leads to long-lasting alterations in the body’s inflammatory response (Danese et al., 2007). Dysregulation of the immune system following chronic stress can result in constant sterile inflammation, which is characterized by increased production of IL-1β and TNFα, in the absence of an immune trigger (Chen and Nuñez, 2010). Such imbalances result in delayed healing (Gouin and Kiecolt-Glaser, 2011) and increased susceptibility to infections and disease progression (Kiecolt-Glaser et al., 1996). As a result, the inflammatory response following an actual immune challenge is expected to be amplified (Frank et al., 2012), causing adverse effects within the brain and body. Lipopolysaccharide (LPS), an endotoxin found in the outer membrane of gram-negative bacteria that are naturally found within the gut microbiome, can result in low-grade inflammation and even sepsis, when it enters the bloodstream (Hasan et al., 2020). An acute injection of LPS is a frequently employed laboratory stimulus to challenge the immune system and such a challenge in a rodent with a history of adolescent stress results in an exaggerated immune response likely mediated by alterations in NF-κB signaling (Bekhbat et al., 2019). NF-κB leads to production of inflammatory cytokines including IL-1β which regulates inflammatory cell recruitment (Gabay et al., 2010) and TNFα which induces fever and inflammation (Dantzer, 2009).

PTSD is characterized by increased anxiety and arousal, intrusive thoughts, and avoidance following life-altering, often chronic, trauma. In addition, PTSD is frequently accompanied by alterations in peripheral system functions (Neigh and Ali, 2016). While it is difficult to model PTSD in rodents, exposure to predatory stress activates similar neural circuitry, including the amygdala, and also precipitates some physiological changes that develop with PTSD (Goswami et al., 2013). In this study, we aimed to determine the extent to which sex impacts the development of anxiety-like behavior and altered peripheral immune response following chronic adolescent stress and to determine if previous reports of increased inflammatory response to LPS following chronic adolescent stress in the hippocampus (Bekhbat et al., 2019) could also be observed in the amygdala.

## Methods

C57BL/6 mice (Male n=16, Female n=16) (Taconic Biosciences, Germantown, NY) were pair-housed with a same sex mate on post-natal day (PND) 22 in ventilated cage racks and were allowed to acclimate for one week before predatory stress. Six Long Evans males (Retired Breeders > 400g), obtained from Charles River Laboratories (Morrisville, NC), were housed individually. Mice were maintained on a reverse 12:12h light:dark cycle, while rats were maintained on a reverse 14:10h light:dark cycle. Although housed separately, both mice and rats received chow and water *ad libitum* in vivaria maintained at 20-23° C and 60% humidity throughout the entirety of the experiment. This experiment was performed in accordance with the Institutional Animal Care and Use Committee (IACUC) of Virginia Commonwealth University, and with the help of the National Institute of Health.

Prior to the start of the predatory stress paradigm, mice were assigned to either the stress (PS) or nonstress (NS) group. PS animals were isolate housed, while NS animals were housed in same-sex pairs for the entirety of the experiment. Mice were placed in a clear 5” diameter hamster ball (Lee’s Aquarium & Pet Products, San Marcos, CA, USA, Cat. #20198), and the lid was secured with 3 alternating pieces of tape, the last of which was covered in crushed rat chow. The hamster ball was placed in the home cage of the Long Evans rat, who was allowed to freely interact with the hamster ball for 30 minutes. The paradigm was conducted for 15 consecutive days from PND 37-51. To prevent habituation, no mouse was paired with the same rat more than 3 times. During this period, NS animals were subjected to daily handling and cage transport. Following each exposure to predatory stress, fecal boli produced by each PS mouse during the stress paradigm were counted and recorded.

Blood was collected via retro-orbital bleed 30 minutes post predatory stress on the 15^th^ day of the predatory stress paradigm. Mice were anesthetized in a small chamber using isoflurane (99.9%/mL) until they no longer produced the toe-pinch response. The blood was initially collected via heparin-coated capillary tubes, then transferred to 1 mL EDTA coated tubes and placed on ice until centrifugation. Blood samples were centrifuged for 20 minutes at 1800g and 4°C, after which the plasma was removed and stored at −80°C until time of assay.

For 3 days prior to behavioral testing, all mice were habituated to the testing suite for 30 minutes. Behavioral assessments occurred on PND 55-56. Locomotor activity and anxiety-like behavior was assessed via the Open Field Task. Mice were placed in the center of the arena (31.75cm x 31.75 cm x 30.48cm) and allowed to roam freely for 10 minutes, before being returned to their home cages. All movement within the open field was recorded and scored by EthoVision XT 14 (Noldus Technologies, Leesburg, VA, USA).

On PND 58, mice were weighed and immediately administered an intraperitoneal injection of lipopolysaccharide (7.5 × 10^5^ EU/kg) from Escherichia coli O111:B4 (Millipore Sigma, Temecula, CA; Cat# L4391). Two hours post-LPS injection, the mice were rapidly decapitated, at which time terminal trunk blood was collected. The blood was processed in the same aforementioned manner. Once extracted, each whole brain was submerged in isopentane for approximately 30 seconds. Amygdala was dissected from each brain, and flash frozen on dry ice until RNA extraction. RNA was extracted using the Qiagen RNeasy MicroKit (Qiagen, Germantown, MD; Cat# 74004). Tissue was homogenized using a 1 mL dounce, and the kit’s provided protocol was used for the remainder of the extraction. RNA was stored at −80°C, until converted to cDNA. The High Capacity cDNA Reverse Transcription kit was used, and samples were diluted in nuclease free water to produce a starting RNA strand of 0.25 μg/ml. Taqman Fast Advanced Mastermix (ThermoFisher Scientific, Waltham, MA; Cat# 4444963) and Taqman Gene Expression Assays (ThermoFisher Scientific) were used. IL-1β (Mm_00434228_m1) and TNFα (Mm_00443258_m1) were normalized against b-Actin (Mm01205647_g1) and GAPDH (Mm_99999915_g1) as housekeeping genes. All LPS animals were normalized against their same-sex control counterparts.

Both baseline and terminal peripheral cytokine concentrations were measured using an MSD VPLEX Proinflammatory Panel 1 Mouse; IL-1β, TNFα Kit (Meso Scale Diagnostics, Rockville, MD, USA)). Peripheral cytokine concentration pooled baseline values are depicted in Table 1. Due to low plasma sample volumes, samples were pooled to produce one baseline sample for each group. Values from each group were averaged to produce a baseline for comparison as depicted in Figure 2.

**Table 1:** Peripheral Cytokine Concentration Pooled Reference Range with Corresponding Coefficients of Variance (n = 1). Values represented are post-predatory stress, but prior to the LPS trigger.

|  | **IL-1β** |  | **TNFα** |  |
| --- | --- | --- | --- | --- |
|  | Concentration<br>(pg/mL) | %CV | Concentration<br>(pg/mL) | %CV |
| **Control Males** | 1.236 | 10.809 | 20.972 | 3.871 |
| **Stress Males** | 1.545 | 3.918 | 16.506 | 0.398 |
| **Control Females** | 1.857 | 6.177 | 23.018 | 4.430 |
| **Stress Females** | 0.654 | 27.188 | 15.227 | 0.610 |

**Figure 2:**
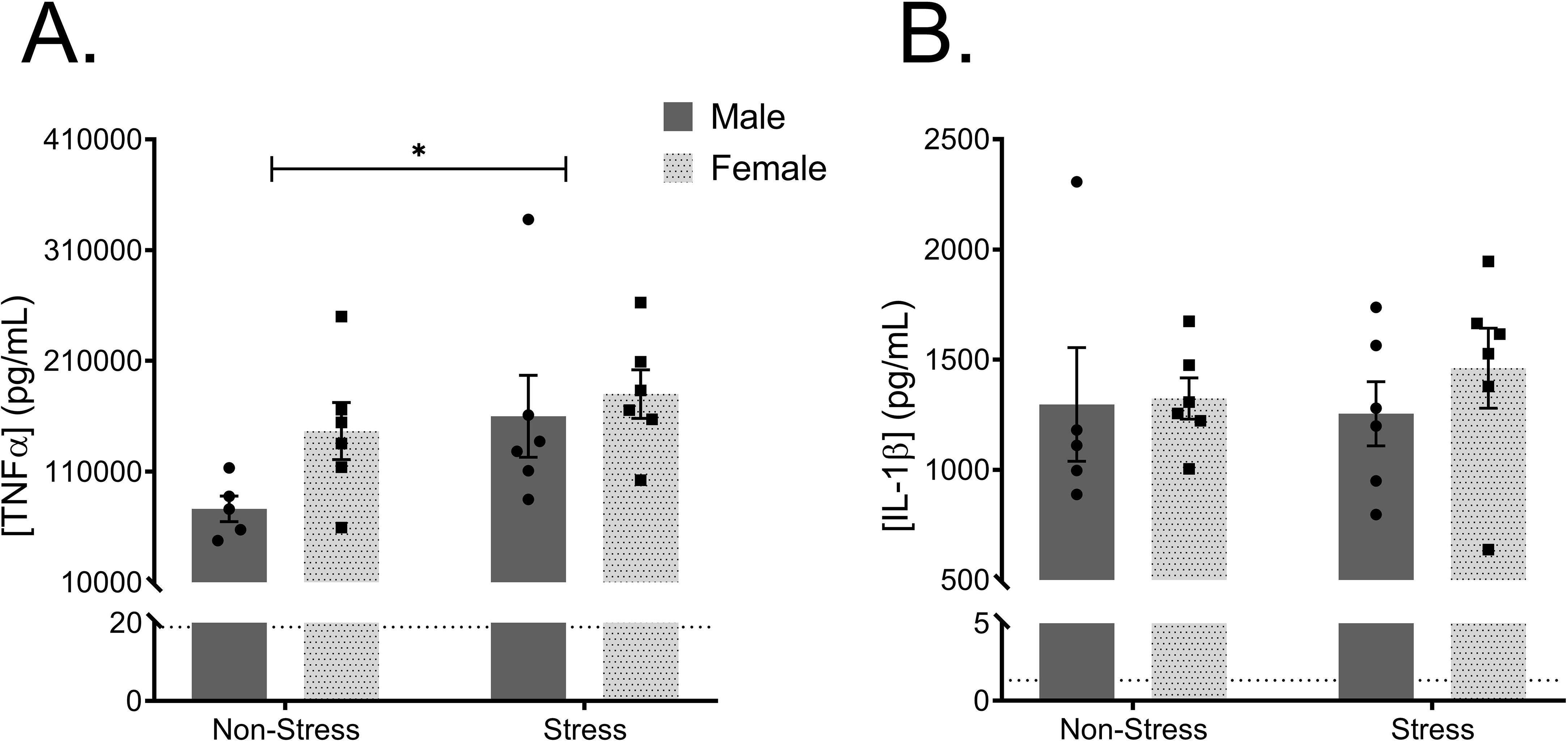
LPS-Induced Peripheral Cytokine Levels. The dotted line on each figure represents the baseline average for each group provided in Table 1. (A) Predatory stress resulted in an increase of circulating TNFα levels (p = 0.0421) following an LPS trigger. (B) However, predatory stress had no impact on circulating IL-1β levels (p = 0.7832) following an LPS trigger.

All statistical analysis was performed in GraphPad Prism v8.3.1 (La Jolla, CA). Biological outliers were excluded following completion of an agglomerative hierarchical analysis using the XLSTAT Excel program (Addinsoft, Inc., 2017). These nine included three NS males, two PS males, two NS females, and two PS females. These samples were excluded from further analysis due to concerns about the adequate delivery of LPS. The α-value was set to 0.05. Two-way analysis of variance (ANOVA) was used to determine the effect of sex and predatory stress exposure on both the behavioral phenotypes following open field and on peripheral cytokine levels. Three-way ANOVA was used to determine the effect of sex, predatory stress exposure, and time on physiological endpoints. Three-way ANOVAs were also used to the determine the effect of sex, predatory stress exposure, and treatment on central expression of cytokines in the amygdala.

## Results

Sex (F_(1, 28)_ = 452.3, p < 0.0001) and age (F_(5.542, 155.2)_ = 8.339, p < 0.0001) dictated group differences in adolescent body mass; however, stress did not significantly alter body mass (F_(1,28)_ = 0.006539, p = 0.9361). As expected, male mice weighed more than female mice, but both sexes gained weight over time. Fecal boli production was significantly influenced by both sex (F_(1, 14)_ = 19.45, p = 0.0006) and length of time in the study (F_(14, 196)_ = 11.29, p = < 0.0001). While females consistently produced more fecal boli during the predatory stress paradigm than males, fecal boli production in both groups decreased over time.

Mice that underwent predatory stress demonstrated an increase in anxiety-like behaviors within the open field (F_(1, 28)_ = 19.82, p = 0.0001), regardless of sex (F_(1, 28)_ = 0.6975, p = 0.4107) as measured by time spent in the center of the arena. Distance travelled and average velocity were also impacted by exposure to stress (F_(1, 28)_ = 43.97, p = < 0.0001), such that mice within the stress group travelled farther and faster than those within the control group. However, following the observation of a stress-sex interaction, post-hoc analysis attributed this difference to a difference between NS and PS females (p < 0.0001).

Circulating IL-1β levels were not affected by stress history (F_(1, 19)_ = 0.07787, p = 0.07787) or sex (F_(1, 19)_ = 0.4607, p = 0.5055). However, circulating TNFα levels increased following predatory stress exposure (F_(1, 19)_ = 4.748, p = 0.0421), but were not impacted by sex (F_(1, 19)_ = 2.817, p = 0.1096). While predatory stress led to changes in peripheral TNFα levels, LPS induction altered both TNFα and IL-1β levels within the brain. TNFα levels within the amygdala, although not affected by sex (F_(1, 31)_ = 0.02275, p = 0.8811) or stress (F_(1, 31)_ = 0.01103, p = 0.9170) history, did significantly increase following LPS induction (F_(1, 31)_ = 51.82, p = <0.0001). Similarly, animals treated with LPS displayed higher levels of IL-1β (F_(1, 31)_ = 38.29, p = <0.0001), regardless of sex (F_(1, 31)_ = 0.06618, p = 0.7987) or stress history (F_(1, 31)_ = 0.2459, p = 0.6235).

The data presented here demonstrate that chronic adolescent predatory stress induces anxiety-like behaviors and exacerbates peripheral inflammatory reactivity, regardless of sex. Exposure to chronic stress during the developmental time of adolescence, leads to immediate consequences including behavioral deficits and inflammation.

## Discussion

Regardless of sex, mice that underwent predatory stress spent less time in the center than their control counterparts (Figure 1). This is indicative of an anxiety-like phenotype within the stress mice. Previous studies have shown that 15 consecutive days of predatory stress can lead to behavioral deficits like those exhibited in this study (Burgado et al., 2014). Additionally, exposure to predatory stress appears to affect locomotor activity, as the stress mice travelled further and with greater speed during the open field task (Figure 1), indicating that stress exposure may lead to hyperactivity (Strekalova et al., 2005). However, post-hoc analysis showed that this effect was driven by the females. During predatory stress exposure, the females did initially produce more fecal boli than the males, however fecal boli production in both groups decreased over time. This suggests that there may be some difference in the colonic activity of males and females, possibly driven by CRH production (Motzer and Hertig, 2004).

**Figure 1:**
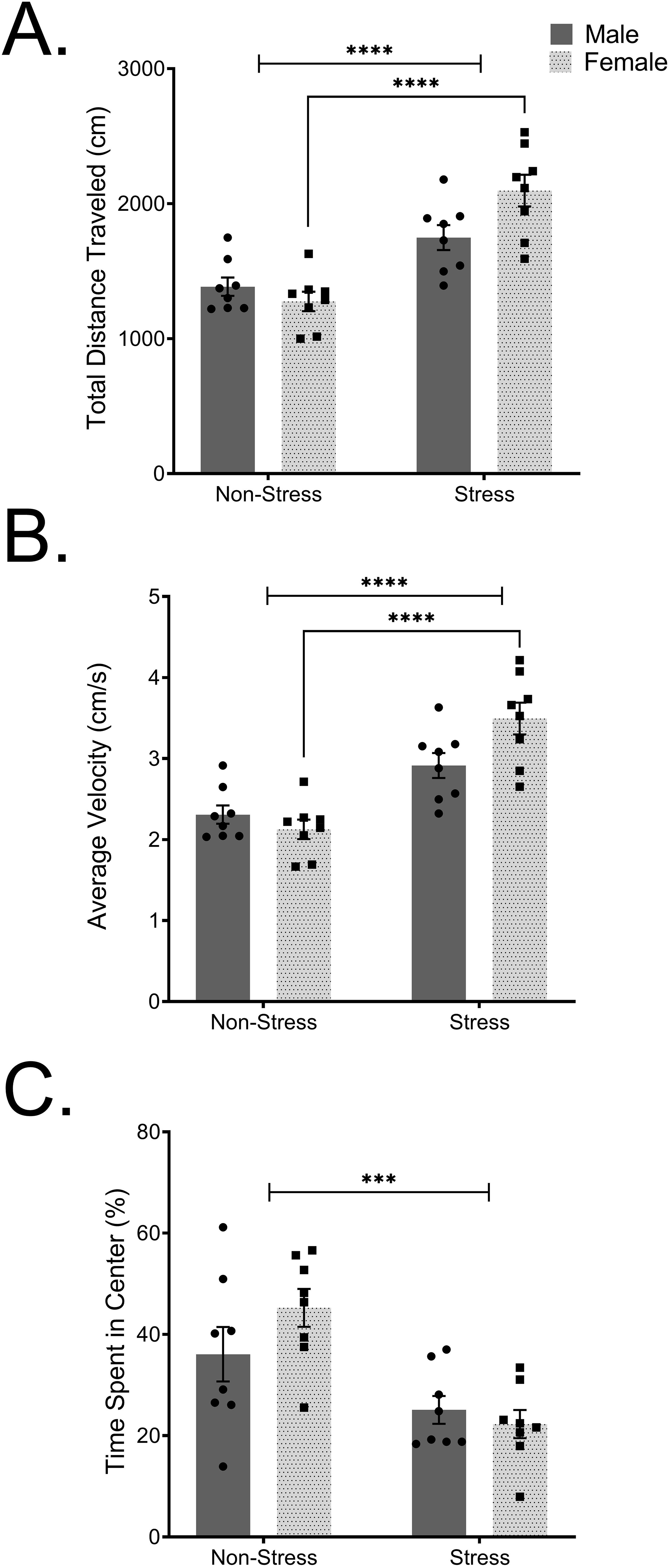
Effect of Predatory Stress on Behavior in the Open Field. (A) Predatory Stress led to an increase in total distance traveled (p < 0.0001) and (B) an increase in average velocity (p < 0.0001). Post-hoc analysis indicated that the main effect of stress on locomotion was driven by the females (p < 0.0001). (C) Control mice spent a greater percent of their time in the center of the field than those in the stressed group (p = 0.0001).

Circulating cytokine levels did not correlate with the observed behavioral deficits as previously suggested by the literature (Bekhbat et al., 2019), but stress-specific changes were observed in the periphery. While IL-1β circulation did not increase following predatory stress exposure in response to an acute inflammatory trigger, peripheral levels of TNFα did, as pictured in Figure 2. In contrast, TNFα and IL-1β within the amygdala, increased in response to LPS to a similar extent regardless of stress exposure (Figure 3). Similarly, Bekhbat et al. (2019) demonstrated exaggerated hippocampal levels of TNFα and IL-1β following LPS induction at the same 2hr timepoint, regardless of sex or stress exposure, in adult rats. However, the impact of stress and sex did not arise until 4hrs post-LPS exposure.

**Figure 3:**
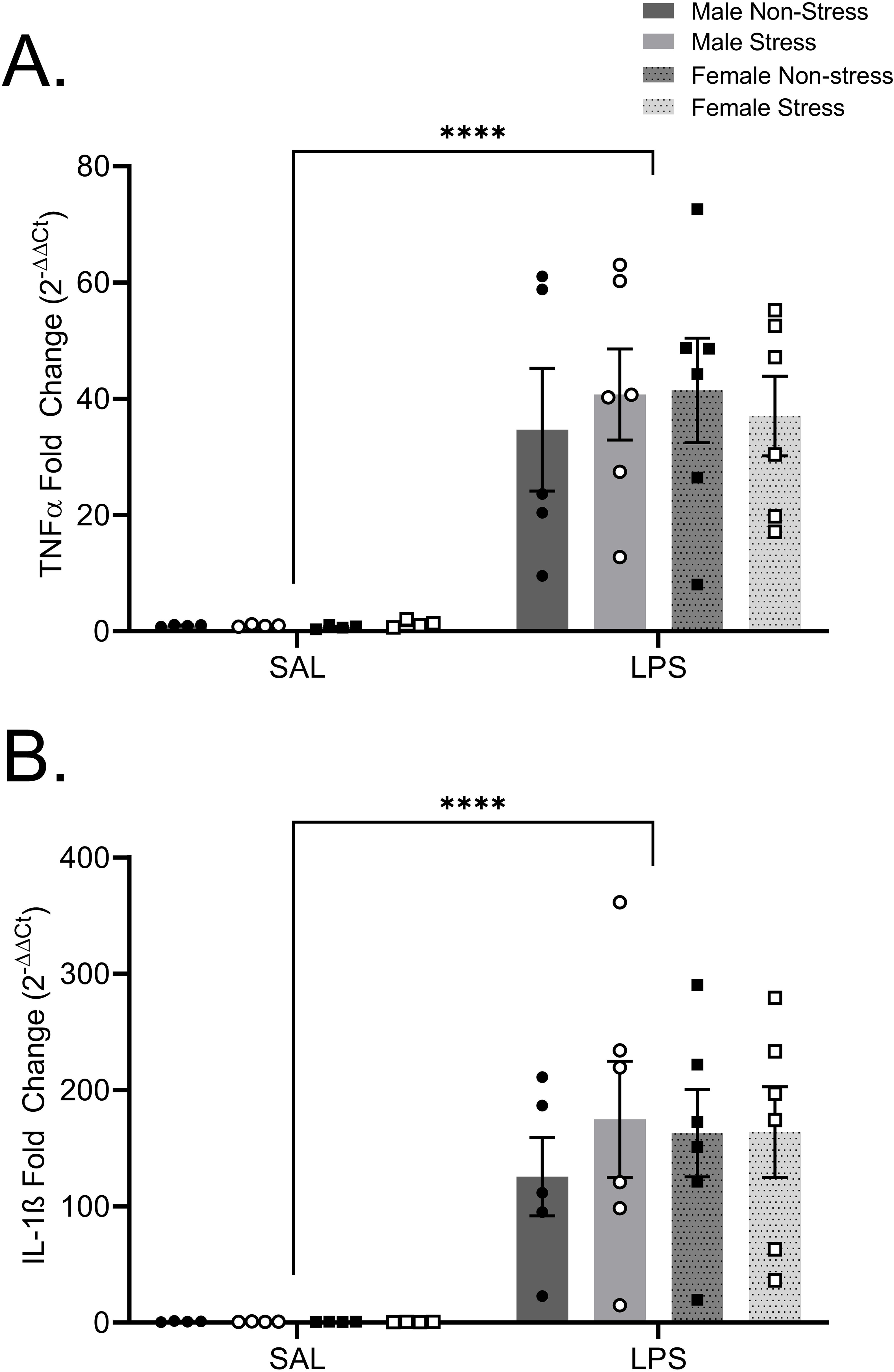
LPS-Induced Central Cytokine Levels. Predatory stress had no impact on (A) TNFα (p = 0.9170) or (B) IL-1β (p = 0.6235) levels within the amygdala. However, there was an increase in both cytokine levels when compared to the saline counterparts (p < 0.0001).

In the current study, we determined that chronic predatory stress exposure has an immediate effect on both the development of anxiety-like behaviors and circulating cytokine levels. The observation that chronic adolescent stress causes an increase in TNFα production following LPS in the periphery, without causing the same response with the brain, suggests that neuronal inflammation is regulated by different mechanics and/or on a different temporal scale. This study did not observe the effects of chronic adolescent stress in adulthood, but previous studies have shown that these behavior and immune alterations persist well into adulthood (Shaw et al., 2020), long after conclusion of the stressor exposure. This study has shown that predatory stress only impacts peripheral cytokine levels, however it is possible that these inflammatory alterations within the brain do not appear until much later, in adulthood or within other brain regions. Additional predatory stress studies are necessary to identify the mechanisms in which stress drives inflammatory alterations, but these data highlight the utility of predatory stress for precipitating similar responses within males and females.

## Acknowledgements

The authors would like to thank the Division of Animal Resources at Virginia Commonwealth University (VCU) for their assistance with animal care and the VCU School of Nursing Biobehavioral Research Laboratory for facilitating completion of the MSD assays. The research was funded by an Alzheimer’s Research Supplement to NINR R014886 (GN).

## Notes

### Competing Interest Statement

The authors have declared no competing interest.

